# PLO(SC)^2^: Plots and Scripts for scRNA-seq analysis

**DOI:** 10.1101/2025.03.09.642205

**Authors:** Markus Joppich

**Author notes:** Corresponding author: Markus Joppich.

## Abstract

**Background:** scRNA-seq analysis has become a standard technique for studying biological systems. As costs decrease, scRNA-seq experiments become increasingly complex. While typical scRNA-seq analysis frameworks provide basic functionality to analyze such data sets, downstream analysis and visualization become a bottleneck. Standard plots are not always suitable to provide specific insight into such complex data sets and should be extended to provide camera-ready, meaningful plots.

**Results:** With PLO(SC)^2^, a collection of plotting and analysis scripts for use in Seurat-based scRNA-seq data analyses is presented, which are accessible for custom script-based analyses or within an R shiny app. The analysis scripts mainly provide a collection of code blocks which enable a comfortable basic analysis of scRNA-seq data from Seurat object creation, filtering, and over data set integration in less than 10 function calls. Subsequently, code blocks for performing differential and enrichment analyses and corresponding visualizations are provided. Finally, several enhanced visualizations are provided, such as the enhanced Heatmap, DotPlot and comparative Box-/Violin plots. These, particularly, allow the user to specify how the shown values should be scaled, allowing the accurate creation of condition-wise plots.

**Conclusion:** With the PLO(SC)^2^ framework data analysis of scRNA-seq experiments is performed more comfortable and stream-lined, while visualizations are enhanced to be suitable for interpreting complex datasets. The PLO(SC)^2^ scripts are available from GitHub and include a vignette showing how PLO(SC)^2^ is applied within a script-based analysis, as well as an R shiny app.

## INTRODUCTION

Single-cell RNA sequencing (scRNA-seq) is becoming increasingly popular. The number of publications on scRNA-seq listed in PubMed nearly doubled from 2020 to 2021 and reached a new record in 2024. As scRNA-seq becomes more accessible and wet-lab protocols become easier to perform, it is also increasingly used to perform large comparisons of multiple scRNA-seq libraries from different disease states. Such complex experimental setups are supported by existing frameworks, but require many function calls. Furthermore, standard plots are not always suitable to provide specific insight into such complex data sets and should be extended to provide camera-ready, fully interpretable plots. This motivates the creation of a collection of scripts to streamline first the pre-processing steps of scRNA-seq experiments, and then the downstream analysis and plotting.

While typical scRNA-seq analysis frameworks such as Seurat (Butler et al., 2018) or scanpy (Wolf et al., 2018) provide functionalities for the analysis of simple data sets, the analysis of complex data sets usually requires additional administrative work by the user: first each library has to be filtered separately, then different libraries have to be integrated, and finally differential gene expression analyses have to be performed for specific conditions. The results are then summarized in various types of plots, such as dimensional plots, dot plots, or heat maps. It is at this stage that visualizations need to be performed with particular care: Dot plots and heat maps often show scaled expression values, so it is important to be able to specify on which data the scaling is calculated.

All of these tasks are included in the PLO(SC)^2^ collection of scripts. For ease of use, the functions included in PLO(SC)^2^ expose only the most important parameters to the user. The functions aim to reduce the amount of code the user has to write and makes it thus easier to handle large data sets. For such large data sets, often consisting of multiple conditions, the default visualizations are not enough to present the data in an adequate way. In order to give an overview of conditions and timepoints, a 2D array of UMAP-plots is useful. Violin-plots are often used to compare values across conditions, and profit from the addition of statistical tests (such as t-test). When comparing multiple conditions or measurement points, data can be visualized side by side. However, when scaled data points are displayed instead of absolute data points, as is often the case in scRNA-seq data analysis, it is important to be able to control which values the scaling was calculated on. PLO(SC)^2^ ‘s enhanced heat map and dot plot make this possible. Much of the functionality of PLO(SC)^2^ is also available through a Shiny app, so PLO(SC)^2^ can be used by more advanced users from scripts, or by anyone using a standalone application.

## IMPLEMENTATION

PLO(SC)^2^ builds upon the Seurat (Butler et al., 2018) scRNA-seq analysis framework and can easily be integrated into new analyses by sourcing the PLO(SC)^2^ scripts from GitHub. It takes advantage of several publicly available R packages for data visualization or analysis, like ggplot2 (Wickham, 2009) and clusterProfiler (Yu and He, 2016; Wu et al., 2021). In general, most functions require a Seurat object as input, as well as a `group.by` and `split.by` clause, identifying the meta-data columns by which plots or analyses are to be generated.

### PLO(SC)^2^ app

The PLO(SC)^2^ app is a graphical user-interface to (most) of the functionality of PLO(SC)^2^. It is implemented using R shiny and converted to a all-in-one executable for the Windows operating system using the R packages *executablePackeR* for creating the portable R environment, and https://github.com/wleepang/DesktopDeployR for creating the stand-alone application.

## RESULTS & DISCUSSION

The PLO(SC)^2^ scripts can be divided into three groups. The first group helps in efficient data processing and provides useful wrappers for Seurat-based (Butler et al., 2018) scRNA-seq analysis. The second group helps in efficient provision of differential gene expression analysis and provides wrappers for clusterProfiler-based (Yu and He, 2016; Wu et al., 2021) gene set enrichment. Finally, the third part of the PLO(SC)^2^ scripts provides camera-ready visualizations for gene expression, like violin- or dot-plots.

### Scripts for Efficient Data Processing

#### Preprocessing data

Each scRNA-seq analysis starts with loading all the data. While the functions provided by Seurat work well for single libraries, all functions must be encapsulated in for-loops or apply-s if these operations are to be performed on many libraries. The wrapper functions provided by PLO(SC)^2^ already operate on a structure suitable for processing multiple related libraries.

Read-in functions are provided for cellranger-generated count matrices, which take a list of h5 or mtx files as input argument (`readH5Files` or `readMtxFiles`). It is automatically checked whether the input matrices contain only expression counts for the RNA assay, or also additional quantification for the antibody capture (e.g. hashtagging or CITE-seq).

The next relevant step is the creation of the Seurat object, which also calculates the mitochondrial and ribosomal RNA fractions (`toObjList` function). Quality control plots and filtering of cells based on their mitochondrial/ribosomal count fraction, the number of features detected and the number of UMIs counted for each cell are performed by the `scatterAndFilter` function. After preparing the RNA assay objects, the antibody capture (if available) can be processed. Within the antibody capture library, hashtag oligo-nucleotides are used to allow the separation of multiple samples per library, or CITE antibodies. For each library, a CITE-seq assay can be added using the `processCITE` function. The user can also define a list of relevant hash tag oligo names for each library. These can be used to identify specific samples within a 10X Genomics library. Processing is performed using the `processHTO` function. The individual libraries can then be split by sample or any other annotated feature, for element-wise integration in the next phase (`splitObjListByGroup`).

#### Integration of Multiple Data Sets

During the integration phase, multiple libraries can be integrated into one Seurat object, e.g. to correct for batch effects introduced during library preparation. Before the actual integration takes place, the list of Seurat objects must be prepared for integration (e.g. identifying common highly variable features) using the `prepareIntegration` function. The integration can then be performed according to the vignettes provided by Seurat on the RNA assay, e.g. using *cca* or *rpca* based integration, or using *SCTransform* processed (Hafemeister and Satija, 2019) data. In addition, joint CITE-seq and RNA-based (multimodal) integration using *wnn* is also possible. The `performIntegration` function collects the most important arguments (e.g. number of PCA dimensions to use) and passes them to the respective Seurat functions.

After integration, the user may be interested in identifying clusters within the combined Seurat object. This can be orchestrated by the `preprocessIntegrated` function, which also allows the user to select the dimension reduction of choice, for example *pca* for normal Seurat objects, *igpca* for the gene expression PCA from *performIntegration*, or *wnn*.*umap* in the case of multimodal integration.

To check whether the integration was successful or whether biases remain, such as cells with few measured features being enriched in one cluster, several quality control plots can be generated and checked with the `makeQCPlots` function.

#### Cell Annotation

Annotating the cells of the Seurat object is one of the most important steps because it defines cell identity in terms of experimental knowledge such as condition, patient, or time point. Within PLO(SC)^2^ the `annotateByCellnamePattern` can be used for this task. It takes as input parameters the Seurat object, the new column name in the metadata, and a list of patterns that describe the new annotation. Each entry in this list of patterns is itself a list defining the new annotation of a cell (e.g. disease or control) and either a set of cell names or a regular expression that can be used to select cells. This selection is also possible on existing metadata columns.

#### Differential Analyses

After filtering cells, performing dimensional reduction, integration, cluster identification and cell type assignment, the key question is often: What are the differences between cell types? To answer this question, marker genes can be calculated for each cluster or cell type. While such functionality is already available in Seurat with the `FindAllMarkers` function, its use is rather indirect: it groups the cells and calculates markers for the currently set global identity. With the PLO(SC)^2^ wrapper, it is possible to specify the groups for which markers are calculated, together with the assay and test to be used, by a metadata column name. The differential gene expression results are annotated with expression data (mean, quartiles, fraction of expressing genes) for both the selected cluster and the background (all other clusters). This information is relevant for the interpretation of fold changes. The whole process is orchestrated by the `makeDEResults` function.

Another common use case is to compare two sets of cells. For this, the `compareClusters` function is provided, which, given two sets of cells, performs a differential gene expression analysis between the two sets and computes gene expression data similarly to the marker genes. Since two sets of cells are often compared for all clusters or any other grouping, the `compareCellsByCluster` function conveniently performs such comparisons. One use case would be to find the difference between each cluster for disease and control samples (e.g. cluster 0 disease vs. cluster 0 control). The results of differential gene expression analysis can be easily visualized as volcano plots using the `makeVolcanos` function. This function takes the list generated by the `compareClusters` function as input and draws a volcano plot for each comparison. The volcano plots are drawn by the EnhancedVolcano package (Blighe et al., 2019). By default, the labeled genes are selected from the EnhancedVolcano library, but the user can specify genes to be shown instead.

#### Enrichment Analyses

The systematic analysis of the enrichment of the gene set is easily done with the `enrichmentAnalysis` extension. To run the enrichment analysis on the Seurat object, you need to specify which Seurat object the analysis should be run on, what the background/universe for set enrichment is, which organism the gene sets should be loaded for (currently human and mouse are supported), and where the gene set enrichment results should be saved (rds file). If all results are to be exported or visualized, an output folder should also be specified. Set enrichment is performed using clusterProfiler (Yu et al., 2012) and ReactomePA (Yu and He, 2016). Both overrepresentation and gene set enrichment analysis are performed on all significantly differentially regulated genes, or only on up- or down-regulated genes. Enrichment analysis on the significantly differentially expressed genes is performed for Reactome pathways (Jassal et al., 2020), GeneOntology (Ashburner et al., 2000), KEGG (Kanehisa et al., 2021) and KEGG modules. In the visualization phase, all result lists are exported to a tab-separated file and Excel, and clusterProfiler’s dotplot, barplot, cnetplot, treeplot and emapplot are also visualized.

### Enhanced Visualizations

After preparing all the relevant data of the scRNA-seq analysis, visualizing these results is (probably) the most time-consuming part of any scRNA-seq data analysis. The plotting functionality provided by PLO(SC)^2^makes the creation of camera-ready plots more convenient.

The PLO(SC)^2^scripts are intended to be used both in a notebook-based environment (such as Rmark-down or jupyter) and in interactive R sessions. For the latter, plots are usually exported directly to png, pdf, and svg formatted files using the `save plot` function. To comply with some journal policies that require submission of source files containing the raw data used for the plots, the data table used by ggplot to generate the plots is also exported to data files (which are tab-separated tables). This way, all relevant files for publishing are created at the time of plot creation.

#### Dimensional Plots

The regular dimension plot (*DimPlot*) in Seurat is used to provide an overview of the preferential 2D embedding of all cells and often provides a first impression of the data, as it is commonly used to show the different cell populations measured in the scRNA-seq experiment. While plotting all cells in this plot provides a broad overview of the data, plotting these dimensional reductions by condition helps to identify condition-specific differences (e.g., the absence of a particular population). The `makeUMAPPlot` function can create dimensionality reduction plots along two axes (e.g., condition and time points, Figure 1). However, different numbers of cells per selected condition can cause a bias in this visualization. Therefore, this function provides the ability to (uniformly) down-sample all conditions so that similar amounts of cells are plotted for each condition. This provides an unbiased view of population prevalence.

**Figure 1.**
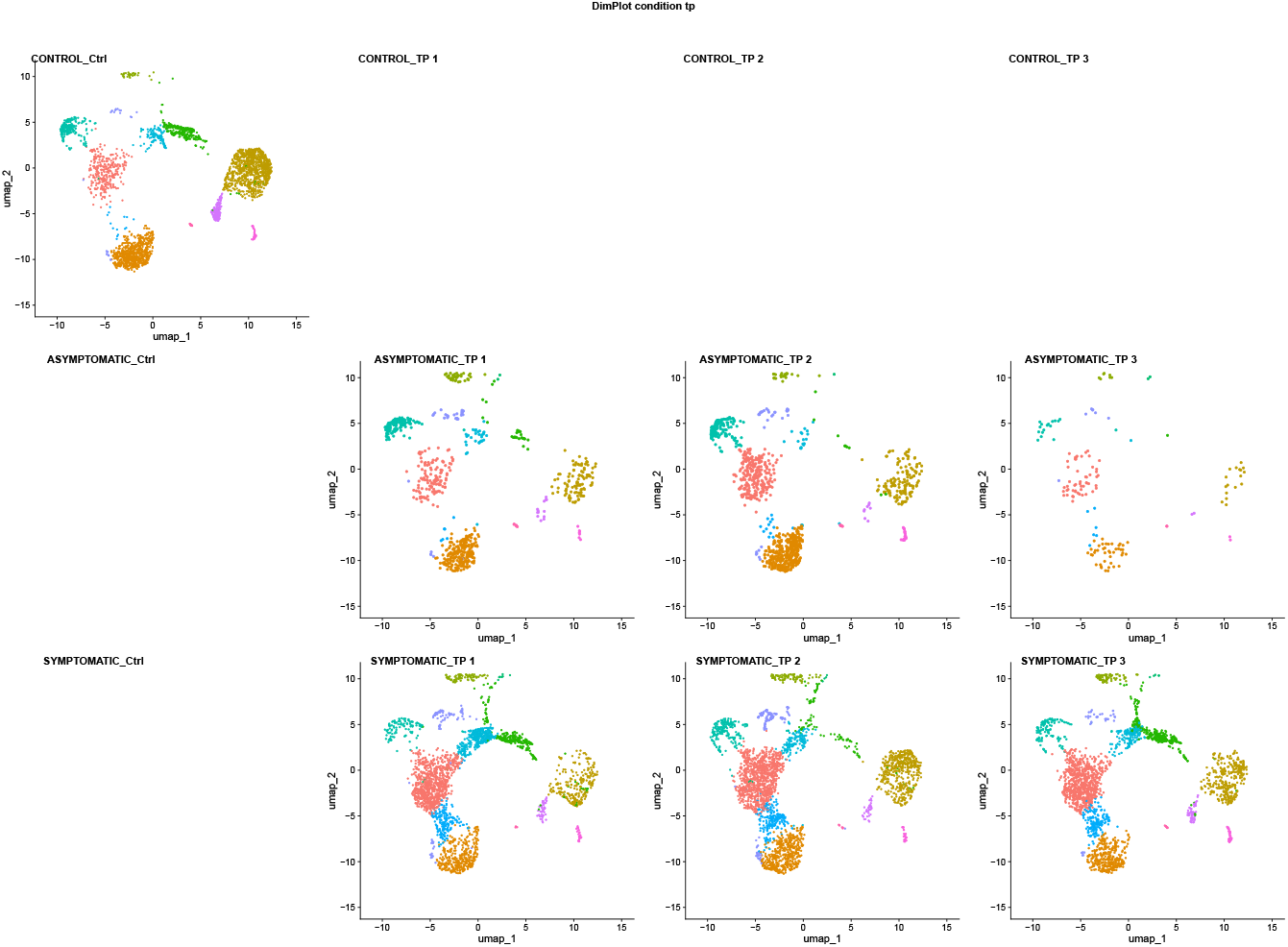
Dimension-wise UMAP-plot. UMAP plot for along multiple dimensions with a shared legend at the bottom. The data set is split along the time series on the x-axis, and the condition (CTRL — ASYMPTOMATIC — SYMPTOMATIC) along the y-axis. While this plot shows all cells, the user can control with a parameter that all sub-plots are down-sampled to the minimal amount of cells within any sub-plot, in order to reduce the visual bias between plots.

Similarly, the expression of certain genes may be compared across levels of a disease (e.g., across multiple time points). While Seurat provides the split.by-option in its FeaturePlot, the individual feature plots have differently scaled legends by default. However, this is not suitable for accurate side-by-side comparisons. To work around this problem, the `splitFeaturePlot` function creates subplots with common legends for all plots, making the individual subplots easily comparable.

#### Violin Plots

The standard violin plot is an important plot type because it shows the expression of a feature over specific groups of the Seurat object. Unlike regular box plots, violin plots also show the distribution of values (Hintze and Nelson, 1998). In general, this is very useful, but it only works as well as the kernel density can be calculated for the violin. Thus, for bimodal distributions or for a small number of values, an additional box plot is beneficial. Therefore, the `SplitVlnBoxPlot` combines both plot types. Also, Violin plots can only be calculated for 3 or more cells. If there are fewer cells for a violin, the violin will not be plotted. This can be a problem when simply overlaying ggplot2’s violin and box plots. Therefore, this implementation can filter out groups with too few cells to make sure that the box plot and the violin plot are well aligned.

Due to the nature of the violin plots described above, these plots are also well suited to visualize two conditions over a time series (Figure 2), where the two conditions are compared for each time point. While the box and violin plots already give an idea of whether there is a change between the two conditions, a t-test can be added to evaluate whether the observed change is statistically significant. This type of visualization is possible with the `comparativeVioBoxPlot` function.

**Figure 2.**
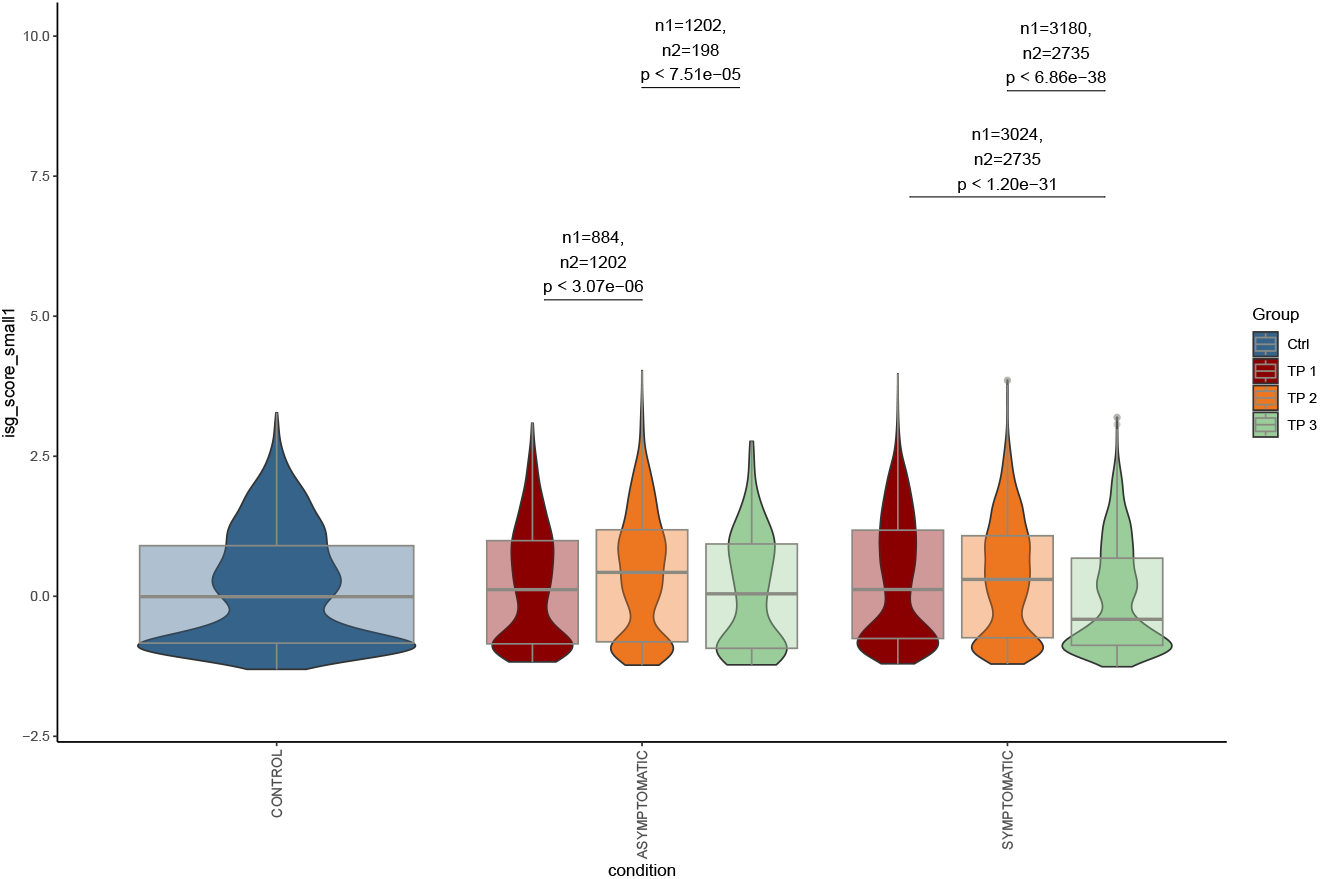
Boxplot and t-test enriched Violin Plot. Violin plots are suitable to show the distribution of the observed values. However, spotting the median from the violin distribution at times can be hard. Here, a regular boxplot becomes helpful. When comparing two conditions for each violin, it might also be interesting to quantify the difference in terms of significance. Therefore, the enhanced violin plot also calculates significance values for the split violins. For this comparison not only the significance value is given, but also the samples sizes which were compared. The user can specify the colors of the groups shown in the plot.

#### Enhanced Heatmap and DotPlot

Heat maps and dot plots are common visualizations in single-cell RNA-seq data analysis. Both are often used to display relative expression values as scaled expression (using z-scores). However, such visualizations must be created with care: Comparing scaled expression values across plots is only valid if both plots have been scaled with the same mean and standard deviation, otherwise the plotted z-scores are not comparable (because they are drawn from different normal distri-butions). In Seurat, the scale.data slot provides a global z-score transformation of all expression values, which is comparable. However, genes may not be part of this slot, or the difference may be too small to be visible. The included advanced heat map and dot plot allow for side-by-side visualizations and provide the option to scale values using different strategies. You can choose to use the globally scaled values from the scale.data slot (`scale.by=“GLOBAL”`), or to scale the gene expression values only for the features included in the plot (`scale.by=“ALL”`). When plots are split by condition, it is also possible to scale the values per subplot (`scale.by=“GROUP”`)). Finally, the raw values of the expression can also be used for visualization (`scale.by=NULL`). This ability to scale the underlying expression values to the user’s needs is the central element of the enhanced plots.

Heat maps are often used to visualize the expression of specific genes across multiple groups of cells, in this case, cell types. While Seurat already provides a *DoHeatmap* function, this heat map shows the expression per cell. While this has the advantage of showing cluster proportions and how many cells express a particular gene, this visualization sometimes becomes too large due to all the information required. A simple heat map then has the advantage of directly showing the expression states per cluster. The advantage of the PLO(SC)^2^ enhanced heat map lies in its usability: the user can specify both the order of genes and the order of clusters, allowing clinicians to visualize exactly the patterns that support their scientific claims. The extended heatmap is accessible via the `makeComplexExprHeatmap` function (figure 3). An extension to this function (`makeComplexExprHeatmapSplit`) allows heat maps to be displayed side by side to compare results from different conditions.

**Figure 3.**
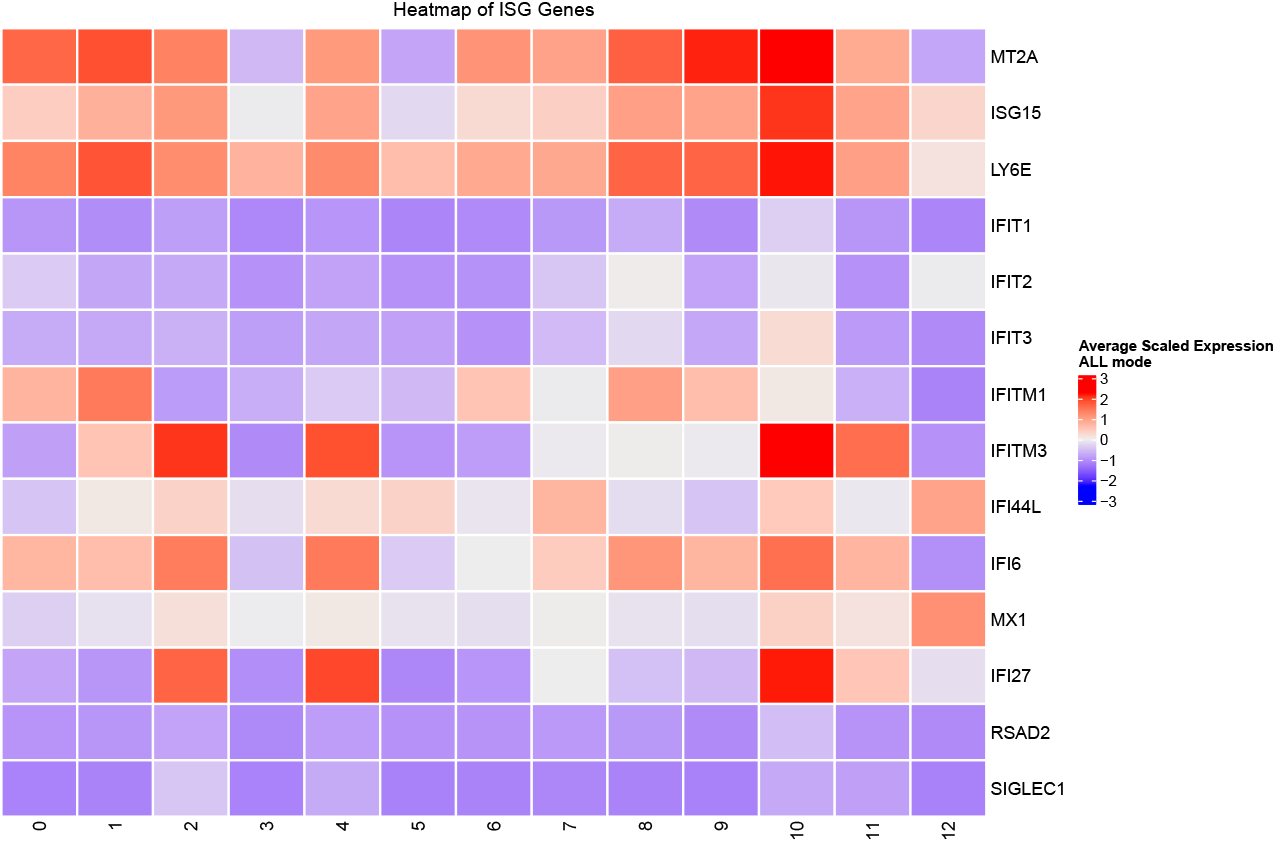
Enhanced heat map of selected genes. Heat maps are commonly used to visualize gene expression of specific genes across several groups of cells - here cell types. While Seurat already provides a *DoHeatmap* function, this heat map shows the expression per cell. This has the advantage of showing cluster proportions and how many cells express a certain gene, this visualization at times becomes too large due to all the required information. A simple heat map then offers the advantage of directly showing expression states per cluster. The advantage of the enhanced heat map is its usage: the user can specify both the order of genes, and the order of clusters, which enables clinicians to visualize exactly the patterns that support their scientific claims. Data can be shown as scaled values either from the Seurat object itself, or z-scaled within all shown values.

Finally, the Dot Plot (`enhancedDotPlot`) is an interesting plot because not only the color of the dot can be interpreted, but also its size. Typically, the color defines the expression strength, while the size of the dot shows the fraction of cells expressing a particular feature. With scRNA-seq data, gene expression is more complicated than with bulk replicates. Gene expression depends on the average intensity of the expressed gene, which depends on the number of cells within a group that express the gene. This is reflected in the dot color and dot size in the DotPlot. In addition, it is also important to know how often a specific group is present within the examined condition: it may be a large group in all conditions compared, or it may be present only in certain conditions. This information is encoded in the background color of the group rows. This addition allows a full evaluation of the expression of a gene in specific groups and between multiple conditions (Figure 4).

**Figure 4.**
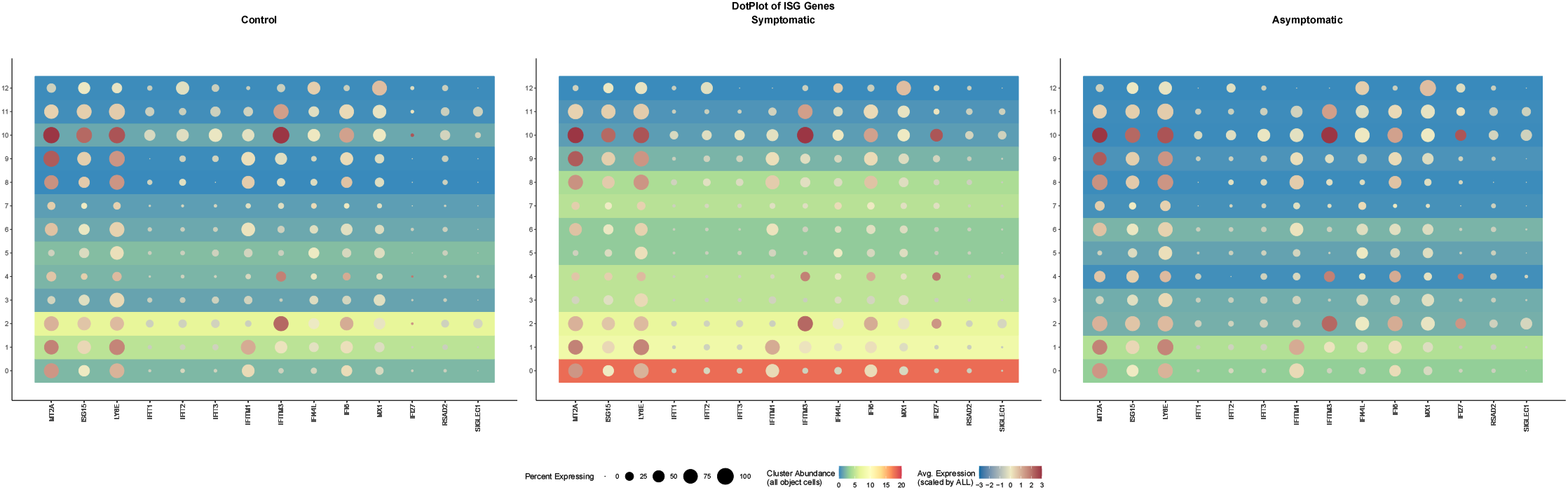
Enhanced Dotplot of selected genes across two conditions. Dotplots are often used in scRNA-seq analysis because they nicely show gene expression per gene and cluster/group, while also showing the percent expressing cells. However, often such information is displayed besides each other, to compare several conditions. Then it must be made sure that the shown data is actually comparable, which is particularly an issue if scaled data is shown. With the enhanced dotplot the user can control on which subset of data the scaled expression should be calculated (e.g. globally, on all shown expression values, etc.). Moreover, the enhanced Dotplot combines gene expression data with group abundance per shown condition (or overall). This allows a complete interpretation of the observed gene expression data.

### Shiny App

In previous work, it was shown that graphical user interfaces can reduce the burden of using bioinformatics software Joppich and Zimmer (2019). In order to facilitate the usage of, primarily, the presented plotting functions, an R shiny app has been developed for PLO(SC)^2^. Using this app, the user can load Seurat objects from an RDS file, calculate differentially expressed genes, perform gene set enrichment analysis and visualize the contained data with the enhanced visualization techniques presented here. Additionally, it is possible to pre-process and integrate new datasets. For ease of use, a stand-alone application of this Shiny app is available for the Windows operating system.

## CONCLUSIONS

The PLO(SC)^2^ scripts are a collection of wrappers for stream-lined data processing and enhanced visualizations in Seurat-based scRNA-seq analysis, helping to make scRNA-seq accessible to beginner and intermediate users. For users proficient with the R programming language, the PLO(SC)^2^ scripts stream-line their scRNA-seq analysis. Instead of writing lines of code for technical purposes (e.g. reading matrices, converting to Seurat objects, writing integration code), the user can focus on the actual tasks. Using quality control plots, it is easy to decide whether a step in the analysis worked satisfactorily or whether other parameters need to be chosen.

The various plot types included in the PLO(SC)^2^ framework allow the creation of professional, informative, and camera-ready plots. With these enhanced plots, it is easy to visualize complex datasets and analyses, while not having to dive into the details of creating these plots on one’s own. In particular, the enhanced HeatMap and DotPlot allow the user to specify which data source to use for scaling the expression data, improving the ability to focus on existing differences. For users who prefer graphical user interfaces, the included Shiny app provides access to most of the features of PLO(SC)^2^and makes scRNA-seq accessible especially for beginners.

The PLO(SC)^2^ scripts and shiny app are available online https://github.com/mjoppich/PLOSC.

## AVAILABILITY AND REQUIREMENTS

Project name: PLO(SC)^2^

Project home page: https://github.com/mjoppich/PLOSC

Operating system(s): Platform independent Programming language: R/Seurat

Other requirements: Seurat 5.0+ (further dependencies, see https://github.com/mjoppich/PLOSC/blob/main/DESCRIPTION)

License: Apache-2.0 license

Any restrictions to use by non-academics: none

## AVAILABILITY OF DATA AND MATERIALS

The PLO(SC)^2^ scripts and Shiny app are available online https://github.com/mjoppich/PLOSC and can easily be installed into R by using `devtools::install github(“mjoppich/PLOSC”)`. PLO(SC)^2^ is distributed under the Apache-2.0 license. The notebook of the analysis of the presented use-case and all required input files are available from Zenodo (https://doi.org/10.5281/zenodo.8268102). The scRNA-seq count matrices are taken from Pekayvaz et al. (Pekayvaz et al., 2022).

## COMPETING INTERESTS

The authors declare that they have no competing interests.

## AUTHOR’S CONTRIBUTIONS

MJ created the PLO(SC)^2^ scripts and wrote the manuscript.

---

**Figure S1.**
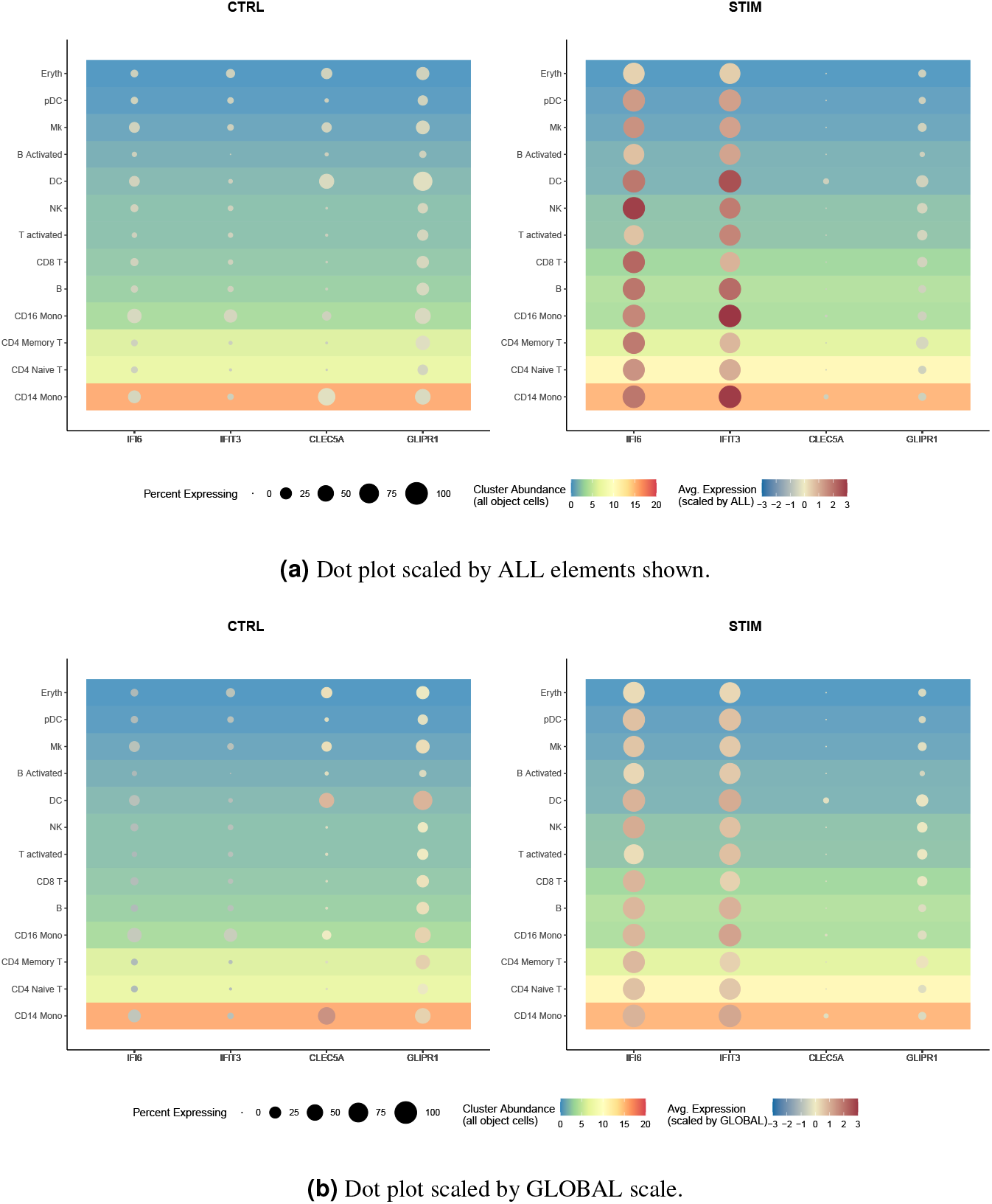
Enhanced dot plot of selected genes in the human IFNB-Stimulated and Control PBMCs dataset taken from the Seurat data library. Values in (a) have been scaled based on all shown values in CTRL and STIM, values in (b) are globally scaled.

## Notes

https://github.com/mjoppich/PLOSC

### Competing Interest Statement

The authors have declared no competing interest.

